# Pinpointing the microbiota of tardigrades: what is really there?

**DOI:** 10.1101/2024.01.24.577024

**Authors:** Bartłomiej Surmacz, Daniel Stec, Monika Prus-Frankowska, Mateusz Buczek, Łukasz Michalczyk, Piotr Łukasik

## Abstract

Microbiota have been proposed as an important aspect of tardigrade biology, but little is known about their diversity and distribution. Here, we attempted to characterize the microbiota of 44 cultured species of tardigrades using 16S rRNA amplicon sequencing, using different specimen pooling strategies, various DNA extraction kits, and multiple types of controls. We also estimated the number of microbes in samples using synthetic DNA spike-ins. Additionally, we reanalyzed data from previous studies.

Our results suggest that the microbial community profiles of cultured tardigrades are dominated by bacterial OTUs and genotypes originating from food, medium, or laboratory reagents. We found microbial strains consistently enriched in certain tardigrades (relative to the culture media and controls), which indicates likely symbiotic associations, but the reads representing putative true tardigrade-associated microbes rarely exceeded 20% of the datasets. Some of the identified tardigrade-associated microbes matched symbionts identified by other studies. However, we also identified serious contamination issues with previous studies of tardigrade microbiome, making some of their conclusions questionable. We conclude that tardigrades are not universally dependent on specialized microbes and highlight the necessary safeguards in future studies of the microbiota of microscopic organisms.

## Introduction

The interactions between eukaryotes and bacteria have become a major topic in modern ecology (e.g. McFall-Ngai et al 2013). Microbial symbionts were shown to influence multiple aspects of the host biology, in many cases having led to important evolutionary transitions. However, the term ‘symbiosis’ encompasses a wide spectrum of fitness effects and degrees of reliance between organisms. Furthermore, despite the ubiquity of eukaryote-bacteria associations, not all eukaryotes are dependent on microbes (Hammer et al 2019). Thus, understanding the microbiota of poorly studied phyla from across the tree of life is critical for the comprehension of the evolution of life on our planet.

Among such ’neglected’ phyla are tardigrades (Tardigrada), also known as water bears. These microscopic animals inhabiting various aquatic and terrestrial environments are best-known for the ability of some species to enter cryptobiosis – a state of suspended animation allowing them to withstand extreme conditions (e.g. Nelson et al. 2015). Recent advances in tardigrade research include the identification of proteins which improve the tolerance to radiation and testing the effects of their expression in human cell cultures (Hashimoto et al 2016), as well as in plants (Kirke et al 2020). However, relations of tardigrades and microbes have been only sporadically investigated, usually by direct observation of bacteria or simple experiments. Examples include the demonstration of the presence of bacteria in specialized vesicles of marine ‘arthrotardigrades’ (Kristensen 1984), discovery of bacteria not being digested in tardigrade gut (Kinchin, 1994), laboratory-induced transport of plant pathogenic bacteria by tardigrades (Benoit et al. 2000), and observations of tardigrade response mechanisms to microsporidial infection (Rost-Roszkowska et al. 2013). These and some other studies were thoroughly reviewed by Vecchi et al. (2016). Interestingly, the abundance of bacteria in tardigrade cultures caused major and well-publicized confusion as the first tardigrade genome assembly was published by Boothby et al. (2015). The authors claimed that an unusually high proportion of genes in the tardigrade genome were acquired by horizontal transfer from bacteria. However, this striking result was rapidly demonstrated to be an artifact resulting from bacterial contamination (Arakawa 2016; Koutsovoulos et al. 2016; Delmont & Eren 2016).

The major breakthrough in microbial symbiosis research was the wide implementation of high-throughput sequencing techniques (HTS), in particular, of amplicons of marker genes such as 16S rRNA, enabling fast and cost-effective screening of microbial diversity across multiple samples. The first attempt to characterize microbial associations of tardigrades using HTS was by Vecchi et al. (2018), who investigated the microbial communities of six tardigrade species. They concluded that the microbiota composition is distinct from the environment and varies between host species, and identified several bacterial OTUs associated with different species of tardigrades. Later, the same samples were used for the analysis of phylogenies and infection rates of potential symbionts (Guidetti et al. 2020). That study also reports the detection of a putative endosymbiont in the ovary of a tardigrade using fluorescence in situ hybridization, suggesting a possibility of vertical transmission of bacteria. Another report on tardigrade microbiota conducted using high-throughput sequencing methods was a conference paper by Mioduchowska et al. (2019) who claimed that tardigrades may host *Wolbachia*. Later, the same team concluded that the microbiota of a newly described species in the genus *Paramacrobiotus* Guidetti, Schill, Bertolani, Dandekar & Wolf, 2009 is partly species-specific (Kaczmarek et al. 2020). Mioduchowska et al. (2023) further investigated *Wolbachia* DNA signal in samples of tardigrades but also crustaceans and molluscs. Arakawa (2020) simultaneously metabarcoded eukaryotes and prokaryotes from moss cushions from wet and dry habitats, demonstrating correlations between abundance of tardigrades and certain bacterial taxa. Relations of tardigrades and bacteria in cryoconite holes were a subject of recent study by Zawierucha et al. (2022), consisting of metabarcoding of bacterial, fungal and eukaryotic genes in samples of tardigrades and substrate. The authors found that microorganisms present in the environment are also present in tardigrades, and spotted significant differences between the microbiota of fully-fed and starved tardigrades. A recent tardigrade microbiota study conducted by Tibbs-Cortes et al. (2022), characterized microbiota of tardigrade communities (tardigrades extracted from samples, without species identification) in American orchards, with the application of decontamination strategy. Using the same samples, the authors observed signs of *Rickettsia* infection in two of 55 tardigrade individuals using FISH (Tibbs-Cortes et al. 2023). Tardigrades were also included in the multi-phylum study of the microbiota of marine invertebrates (Boscaro et al. 2022), which revealed no clear correlation between the microbial community composition and the host phylogeny.

Unfortunately, the primary method used for these studies, marker gene amplicon sequencing, has multiple biases and caveats that need to be addressed carefully before the results can be considered reliable (Knight et al. 2018). Among these problems is contamination, to which low-biomass samples such as tardigrades are particularly prone, and which may dramatically affect the results (Eisenhofer et al. 2019, Salter et al 2014, Hammer et al. 2019). Contamination categories include, but are not limited to (1) environmental contamination, where samples may contain bacteria originating from food or living in culture medium, but not specifically associated with the host organisms; (2) contamination from reagents, including DNA extraction kits and PCR mixes (Hornung et al. 2019), shown to strongly affect the microbiota profiles in some low-biomass samples (Salter et al. 2014), (3) cross-contamination during library preparation & sequencing – in multiplexed Illumina flow cells, reads from one sample may be incorrectly assigned to another (index hopping) (van der Valk et al. 2020, Illumina 2021); (4) analytical challenges, where comparisons based on often imprecise taxonomic assignments obscure biological patterns and make it hard to detect the issues listed above. On top of these methodological challenges, the microbiota of micrometazoans can be highly variable within and across species, populations, environmental conditions or sampling times (Eckert et al. 2020), which together with contamination issues, may substantially complicate and confound the conclusions. Therefore, a careful selection of controls, replicates and other methodological approaches is of crucial importance when studying microbiota compositions in low-biomass samples. Replicate studies and reanalysis of the already published datasets might be necessary to obtain an accurate picture of microbial associations in organisms such as tardigrades.

In this study, through a series of carefully designed sequencing-based experiments, we aimed to identify bacteria truly associated with tardigrades, while controlling for several other sources of microbial signal. After our first surveys had highlighted strong experimental noise (contamination from different sources), we tested different sample preparation methods using multiple cultured tardigrade species and examined various negative samples representing different components of the laboratory environment to pinpoint the real associations between tardigrades and bacteria. Next, we carefully assessed and controlled various sources of experimental noise when characterizing the microbiota in 12 cultured tardigrade species. Furthermore, we critically re-analyzed amplicon data from previous studies on tardigrade microbiota. We show that the vast majority of bacterial signal in tardigrade microbial community profiles, whether sequenced by us or other authors, originates from sources other than the tardigrades themselves. Based on these results, we provide recommendations for future studies of microbial communities of tardigrades and other microscopic eukaryotes.

## Material and methods

### Study design

This study comprised five parts (Fig. 1). We started from (*i*) a broad survey of microbiota across multiple tardigrade cultures representing different species, and (*ii*) a more systematic effort to compare replicate cultures of three tardigrade species. After realizing the extent of microbiota overlap across experimental and negative control samples, we endeavored to systematically assess sample processing methods and methodological caveats before addressing biological questions. Hence, we (*iii*) compared different tardigrade sample pooling and processing approaches. We then attempted (*iv*) a careful microbiota comparison of selected tardigrade cultures. Finally, we (*v*) reanalyzed data from previous studies on tardigrade microbiota and compared the results with our dataset.

**Figure 1.**
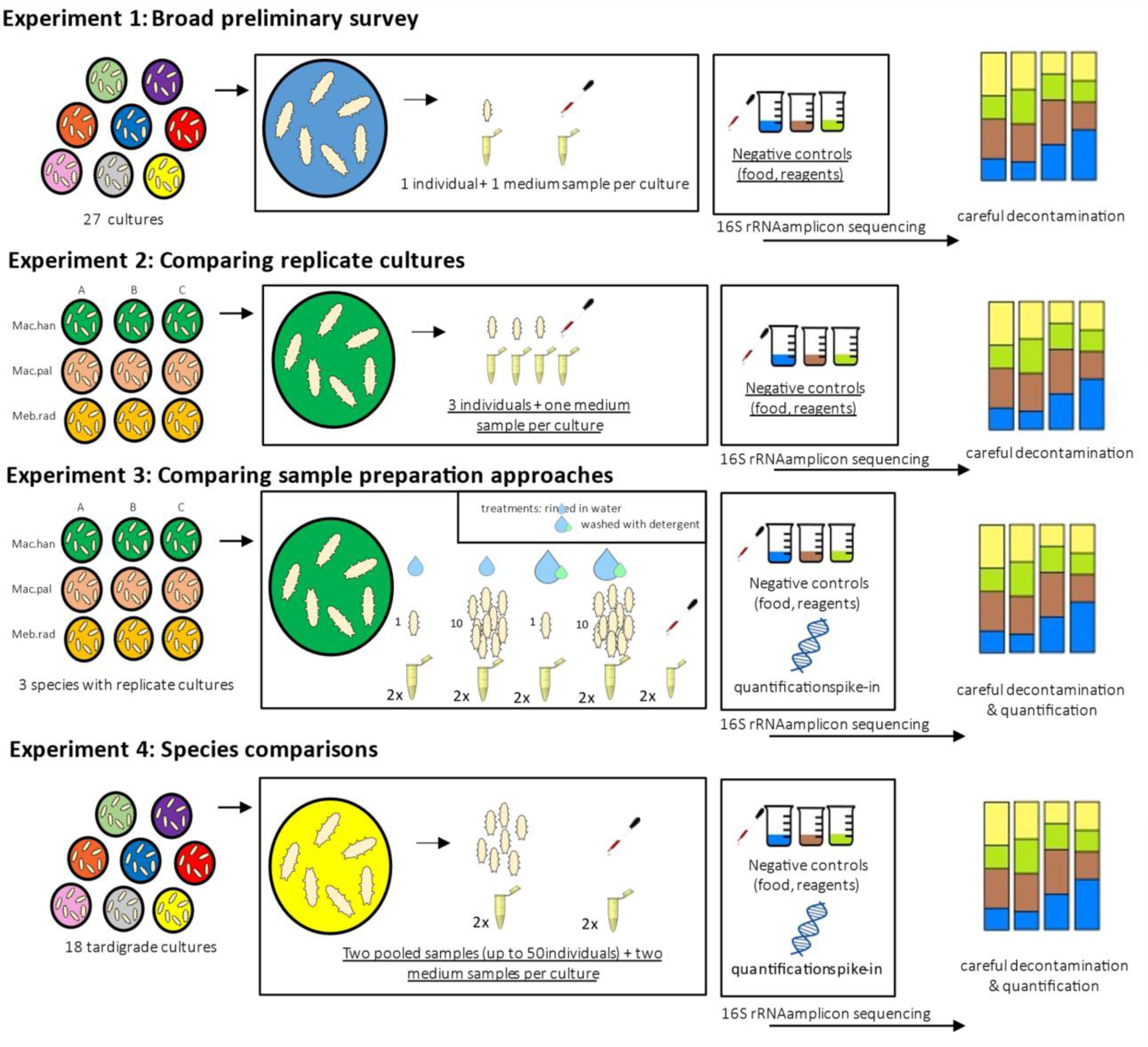
A visual summary of experimental approaches used in this study.

### Study material

All experimental procedures were conducted using 44 tardigrade cultures maintained at the Institute of Zoology and Biomedical Research of Jagiellonian University and representing substantial phylogenetic diversity: 38 species representing seven genera of the families Macrobiotidae (30 spp.), Hypsibiidae (4 spp.) and Milnesiidae (4 spp.); see Table S1 for details. The analyzed cultures were established between 2013 and 2019 from at least several live individuals extracted from a single substrate sample (moss or lichen cushions), and maintained in the laboratory since collection. Additionally, the dataset included a culture of *Milnesium inceptum* established by Ralph Schill in 2003 and a commercially available strain of *Hypsibius exemplaris* from Sciento, originally established in 1987 by Robert McNuff.

Cultures were kept and maintained under standard laboratory conditions in Petri dishes placed in climatic chambers set to 16 °C and complete darkness (Stec et al. 2015, Kosztyła et al., 2016) in a medium consisting of ‘Żywiec Zdrój’ spring water. Once a week, all cultures were supplied *ad libitum* with food (rotifers *Lecane inermis* Bryce, 1892 and/or algae *Chlorella* sp., both from long-term laboratory stocks). Once per month, medium within each tardigrade culture Petri dish was vigorously mixed and after quick sedimentation of animals and eggs, old medium was collected using a glass pipette under stereomicroscope and discarded. Then, the cultures were refilled with clean spring water and fed as described above.

### Experiment 1: Broad preliminary survey

The first experiment was designed to test the suitability of methodological approaches for addressing questions of how distinct and consistent are the microbiota of different tardigrade species. We asked whether single tardigrades provide a strong microbial signal, different from that in the culture environment and food. In this experiment, we used a single tardigrade individual and a 100 μl culturing medium sample from each of 30 cultured species (Table S1). DNA was extracted using the Chelex 100 resin (Bio-Rad) extraction method (Casquet et al. 2012) with modification described in Stec et al. (2020). Along with samples of tardigrades and culture media, samples of tardigrade food (rotifers *Lecane inermis* and algae *Chlorella* sp.) were prepared by pipetting 100 μl of their main stocks, each in two replicates. Additionally, a DNA extraction blank and three PCR blanks were prepared.

### Experiment 2: Comparing replicate cultures

In the second experiment, we tested whether replicate individuals and subcultures of each species have consistent microbiota that differ significantly among species. We addressed these questions using replicate individuals and medium samples from different laboratory stocks of three tardigrade species. The cultures of three selected tardigrade species: *Mesobiotus radiatus* (Culture ID: KE.008), *Macrobiotus hannae* (PL.010) and *Macrobiotus ripperi* (PL.015) consisted of three replicate stocks – subcultures that were kept separated for at least 2 years before the experiment (although provided with the same medium and food, providing possible routes of microbial transfer). Four samples (three individual tardigrades, rinsed several times with sterile water, and a 100 μl sample of culturing medium) were taken from each of the three subculture dishes of each species, for each of the two DNA extraction methods (Fig. 1). The first was a Chelex 100 resin (Bio-Rad), as described above, and the second a custom method combining bead beating of tardigrade samples suspended in 200 ul of lysis buffer (0.4 M NaCl, 10 mM Tris-HCl, 2 mM EDTA, 2% SDS, 1.0 mg/ml Proteinase K) on Omni Bead Ruptor Elite homogenizer (twice for 30 s with 5m/s speed), followed by 2 h digestion at 55oC, and lysate purification using Sera-Mag SpeedBeads (SPRI) (Rohland & Reich 2012). Regarding negative controls, we used extraction, PCR, and food controls for the Chelex method as listed in Experiment 1, and analogous control for the SPRI bead method.

### Experiment 3: Comparing sample preparation approaches

In order to improve the signal-to-noise ratios in microbiota reconstructions, we decided to compare alternative sample pooling, their subsequent treatment, and DNA purification strategies more systematically. We used the same nine subcultures of three species as in Experiment 2. The tardigrades extracted from Petri dishes were subjected to one of the two treatments: (*i*) rinsing in sterile water (*ii*) rinsing in sterile water with addition of dish soap ‘Ludwik’ to aid the removal of topical microbiota from tardigrade cuticle. For each subculture, for each of the two treatments, we prepared five total samples: two individual tardigrade samples, two pooled samples of 10 animals, and one 100 µl sample of culture medium. DNA was extracted using the custom method described above. Additionally, for the three cultures of *Macrobiotus ripperi* (PL.015), the procedure was repeated using another DNA extraction method – DNeasy Blood & Tissue Kit (Qiagen Ltd.), a standard kit in invertebrate microbiota studies. Negative controls of algae, rotifers, ’Żywiec Zdrój’ spring water, water with detergent, and extraction blanks were prepared in two replicates each, for each of the two DNA extraction approaches. To allow amplicon quantification after DNA sequencing, we used quantification spike-in plasmids containing an artificial sequence flanked by conserved 16S rRNA priming sites, Ec5502 (Tourlousse *et al*. 2017). To one-fifth volume of each homogenate sample we added ca. 1000 copies of the plasmid. In samples processed by Qiagen extraction method, the same number of copies was added, but to the whole extract volume.

### Experiment 4: Species comparisons

For tardigrades from 18 cultures of 12 species, including all nine cultures of three species used in Experiments 2–3, four cultures used in Experiment 1, and five cultures not included in the previous experiments, we prepared two pools of 5–50 individuals, depending on availability (Fig. 1; details in table S1). For each culture, we also used two culturing medium samples, and also included in the experiment the full range of controls, as listed above. We processed the samples using methods validated during Experiment 3: DNA was extracted by SPRI Beads method and an aliquot of approximately 1000 copies of quantification spike-in standard Ec5502 was added to one-fifth of the homogenate volume for each sample prior to purification.

### Amplicon library preparation & sequencing

We prepared amplicon libraries for the V4 hypervariable region of the bacterial 16S rRNA gene using a custom two-step PCR approach, as described recently (Michalik et al. 2021). In the first round of PCR, we amplified two marker regions of interest using template-specific primers 515F/806R (Caporaso et al. 2010), with variable-length inserts and partial Illumina adapters. In the same multiplex PCR reactions, we amplified other targets, but these data were not used. The SPRI bead-purified PCR products were used as templates for the second, indexing PCR. Both the first and the second PCR reaction were conducted in 10 μl volumes, including 5 μl of Qiagen Multiplex PCR Mastermix and primers at the concentration of 1 μM, and cycling conditions that included 27 and 10 cycles, respectively.

Libraries were pooled based on band brightness on an agarose gel, and pools sequenced across three multiplexed Illumina MiSeq v3 lanes (2×300bp reads) at the Institute of Environmental Sciences of the Jagiellonian University.

### Data analysis

#### Data preprocessing

The raw reads were processed following a modified pipeline of the Symbiosis Evolution Group at the JU, and the procedure with detailed explanation of each step is stored at a repository (github.com/bsurmacz/Tardigrade_microbiome). Briefly, paired reads were assembled into contigs and quality-filtered using PEAR v0.9.11 (Zhang *et al*. 2014), then dereplicated, denoised and screened for chimeras using USEARCH-UNOISE3 (Edgar 2010). The resulting denoised zero-radius Operational Taxonomic Units (zOTUs) were clustered into OTUs with a 97% identity threshold. The zOTUs as well as the OTUs were classified taxonomically using VSEARCH (Rognes *et al*. 2016) through a comparison against the SILVA SSU 138 database (Quast et al. 2013). The classification was done for the merged dataset from all the experiments to allow uniform (z)OTU names in the study. All pipelines and scripts used for these steps are available from the github repository listed above.

#### Contamination filtering, classification, and comparison

The custom Decontamination step was done using data from each experiment separately (except for Experiments 1 and 2, where samples were processed at the same time, we used overlapping negative control samples). Identification of contaminants in samples was performed using a custom script (github.com/bsurmacz/Tardigrade_microbiome), expanding upon a workflow developed and implemented previously (Łukasik et al. 2017). As an input, we used a zOTU table for all samples from an experiment, including negative controls (DNA extraction blanks, PCR blanks), tardigrade food (algae, rotifers), culture medium samples, and tardigrades. For each zOTU, we computed its maximum relative abundance (MaxRelAbund) in each sample type. By comparing the values across sample types, we assigned each zOTU to one of six categories: PCR contaminants, extraction contaminants, rotifer-associated microbes, algae-associated microbes, spike-in reads, and tardigrade symbionts.

Starting at the beginning of the above list of categories, we looked at control samples for the given category, reasoning that zOTUs that are not substantially more abundant in samples other than these controls should be classified as representing the given category. For example, we assumed that a zOTU that is present in some PCR negative controls, and not substantially more abundant in at least some samples other than these controls, should be classified as a PCR contaminant. As a “substantially more abundant” threshold we have set a value of 10, identified previously as providing a good balance between sensitivity and specificity and verified experimentally (by comparing results of decontamination using different thresholds); Łukasik et al. (2017). Thus, we started from classifying as PCR contaminants all zOTUs with MaxRelAbund in samples other than PCR blanks less than 10x MaxRelAbund in PCR blanks. Subsequently, after removing from the dataset all PCR blanks and PCR contaminant zOTUs, we did the same comparisons for DNA extraction blanks, and then again for algae and rotifers. Finally, using the same method, we indicated bacteria enriched in tardigrades relative to culture medium. Reads representing the artificial spike-in target Ec5502, added to all samples in Experiments 2 & 3, were not removed, but, instead, they were used for the template quantity estimation, as explained below. The estimated proportions of contaminants in different categories were analyzed in a full-factorial design, employing extraction method, pool size, treatment (cleaning), species, and culture replicates (Supporting Information 2). The effects of above mentioned factors were tested by constructing beta-distributed mixed-effects generalized linear models, implemented in R package ’glmmTMB’ (Magnusson et al. 2017) with logit link and beta distribution with culture replication used as a random factor. Model diagnosis was made by visual inspection of residuals from the DHARMa package (Hartig 2022). We used a similar approach to test the effects of the factors on microbiota community composition. To achieve this, we applied PERMANOVA using the *adonis* function in the ’vegan’ R package (Oksanen et al. 2022). The calculations were done using R.4.1.3 (R Core Team 2013).

We estimated the starting amount of 16S rRNA copies in the processed samples – tardigrades, culture medium, but also food and control samples in Experiments 3 and 4 – based on the proportion of reads representing quantification spike-in plasmid Ec5502 (Tourlousse et al. 2017). As stated before, known amounts of copies of the linearized plasmid carrying artificial targets amplifiable with 515F-806R primers had been added to homogenized samples prior to purification. Assuming no strong bias for or against the artificial target (relative to bacterial 16S rRNA) during purification, PCRs, or sequencing, we anticipated that the ratio of plasmid to symbiont signal will reflect the starting amount of 16S rRNA copies after homogenization (Tourlousse et al. 2017). Hence, we computed the starting 16S rRNA copy numbers by multiplying symbiont/spike-in read ratios by the number of plasmids added (1000 copies). For the SPRI bead-purified sample, the resulting value was further multiplied by five, as only one fifth of the available homogenate was mixed with 1000 plasmids.

#### Reanalysis of the data from previous studies

We identified two 16S rRNA amplicon sequencing datasets representing studies that attempted to characterize and compare tardigrade microbiota, but did not utilize robust contamination controls. First, we downloaded from NCBI short read archive data from three studies by a Polish consortium: Mioduchowska et al. (2019) PRJNA530068; Kaczmarek et al. (2020) PRJNA549029; Mioduchowska et al. (2021) PRJNA758403). Simultaneously, we downloaded data from a microbiota study from the same laboratory but which focused on mussels (Mioduchowska et al. 2020), which we thought might allow the identification of potential common laboratory contaminants. All the above mentioned libraries targeted the V3-V4 region of the 16S rRNA gene. In total, we downloaded 10 libraries for tardigrades, 3 libraries for the tardigrade culture environment (rotifers and rotifer medium) and 19 libraries for other organisms. The second dataset consisted of a single study (Vecchi et al. 2018) PRJDB5471. In this study, V4 region of 16S rRNA was amplified from tardigrade and tardigrade environment samples and two negative controls. These two datasets were analyzed separately, following the same pipeline as described above. For contaminant identification in the Vecchi et al. (2018) dataset, we used their two negative controls (joint for the DNA extraction and the PCR step). For the former dataset, with no negative controls provided, we used proxy libraries from the same group’s mussel microbiota study as negative controls. We compared the taxonomic annotations and sequences of abundant OTUs in the reanalyzed datasets with our data.

## Results

### Data summary, and the diversity of sources of microbial signal

Across the four experiments, we obtained a total of 7,919,701 reads. Of these, ca. 58% reads on average had error-free primers corresponding to the V4 region of 16S rRNA, and only these reads were used for subsequent steps; we did not use the remaining reads containing errors or representing other regions that were co-targeted in the same PCR reactions. After discarding libraries with very low numbers of quality 16S-V4 reads (<100) and libraries of culture medium whose tardigrade samples did not pass the threshold, we retained data for 196 tardigrade samples (pooled or individual), 92 samples of tardigrade culture medium, and 47 samples representing four types of negative controls: PCR blanks, extraction blanks, algal cultures, and rotifer cultures. Numbers of reads for these different categories are provided in Table S2.

Initially, we focused analyzes on the microbial community profiles of control samples, attempting to identify the dominant sources of contamination in tardigrade samples. We found that each of these four categories had a consistent, characteristic microbial community profile (Fig 2A). PCR blank profiles were dominated by the signal of *Brachybacterium* sp. (which we previously identified as a contaminant in Qiagen Multiplex PCR Mastermix). Other common taxa classified as PCR contamination were *Pseudomonas* and *Methylobacterium*-*Methylorubrum*. Extraction contaminants were more diverse, and more variable among methods and sample batches. The most common bacterium in that category was *Lactococcus*, abundant in Chelex extraction blanks, but also found in samples extracted using the SPRI bead method. Algae libraries contained a high proportion of non-bacterial reads – up to 29% of reads represented mitochondrial 16S rRNA of *Chlorella*. The microbiota associated with rotifer cultures was more diverse, with most abundant bacteria classified as *Legionella*, *Polynucleobacter*, *Microbacteriaceae* and *Comamonadaceae* (Fig 2A).

**Figure 2.**
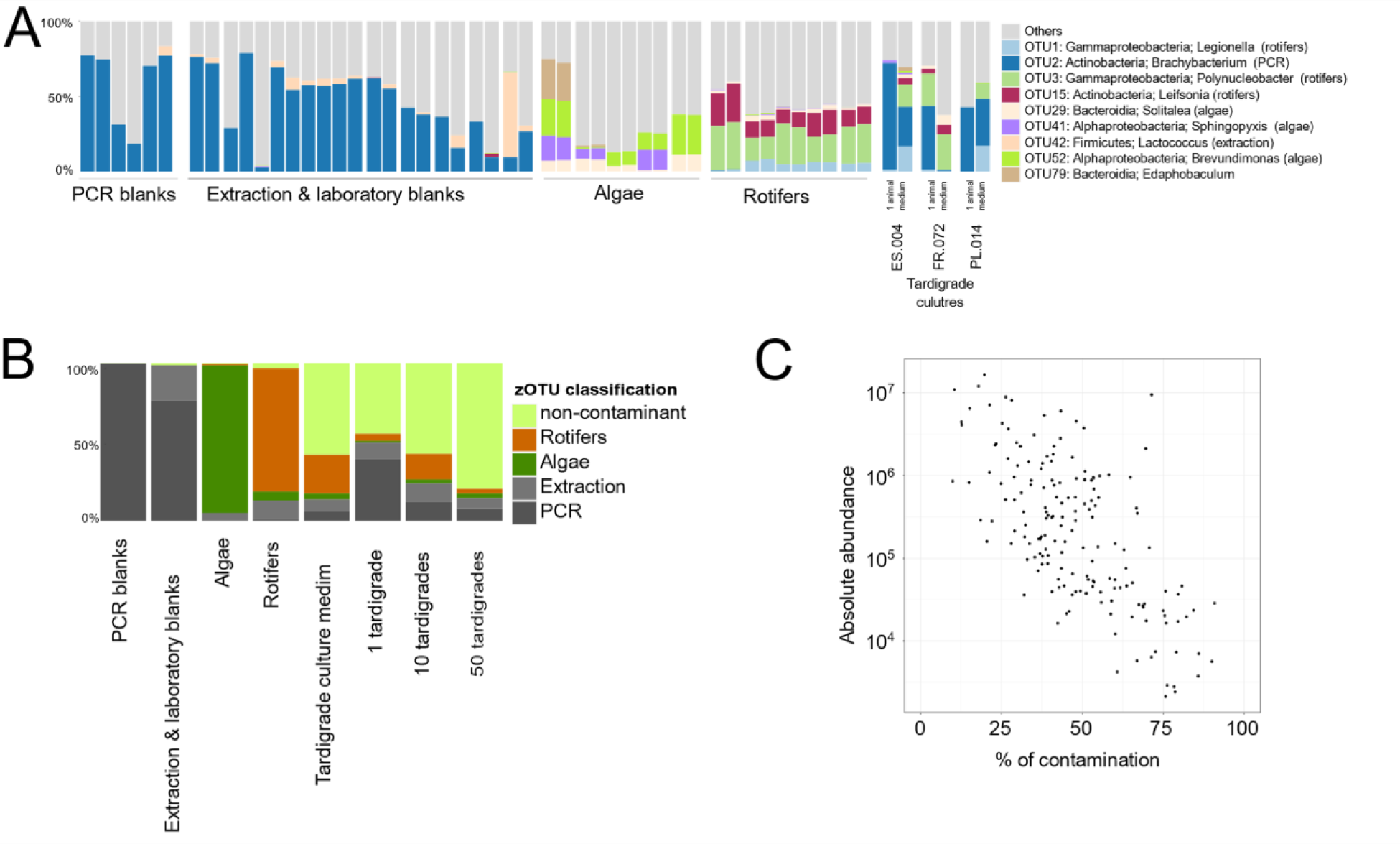
Overview of the contamination present in the dataset: **A** – distribution of most common OTUs in various types of blanks compared to representative tardigrade and culture medium samples from across Experiments; **B** - average percentage of tracked contamination sources in different types of samples across the study; **C** – relation between estimated absolute abundance of 16S rRNA copies in tardigrade and medium samples (after removal of DNA extraction and PCR contaminants) in samples of tardigrades or tardigrade culture medium from Experiments 3 & 4.

Bacteria dominant in these four types of controls comprised a large portion of the microbial community profiles in tardigrade and tardigrade medium (Fig 2B). In Experiments 1 and 2 (with a single tardigrade, or a drop of medium, used as starting material), libraries contained up to 92% of contaminant reads (Table 1), with PCR contamination making up about three-quarters of the experimental noise (Table 1, Figure 2B). In comparison to tardigrade samples, culture media had a 5-fold lower extent of PCR contamination but much higher share of contaminants from rotifers and algae (Table 1).

**Table 1.** Relative abundance of different types of contamination identified in tardigrade and medium samples in different experiments. For each category of samples and each type of contamination, we provided median (and min-max) values.

| Sample | PCR | Extraction | Algae | Rotifers | Non-contaminants | Absolute abundance of 16S rRNA copies |
| --- | --- | --- | --- | --- | --- | --- |
| **Experiments 1 & 2** |  |  |  |  |  |  |
| Tardigrades (1) | 34.5% (0.7–91.9) | 6.01% (0.0–31.3) | 0.52% (0.0–8.4) | 1.2% (0.0–35.1) | 51.6% (8.1–99.0) | – |
| Medium (100 µl) | 7.2% (0.0–44.6) | 3.95% (0.0–21.3) | 3.2% (0.7–30.3) | 19.0% (0.0–62.5) | 62.2% (13.8–92.6) | – |
| **Experiment 3** |  |  |  |  |  |  |
| Tardigrades (1) | 35.8% (21.4–71.1) | 17.3% (7.2–26.4) | 0.7% (0.0–8.5) | 8.0% (0.0–22.3) | 30.8% (14.1–57.5) | 17,845 (1,552–266,250) |
| Tardigrades – washed (1) | 40.0% (17.4–69.1) | 13.7% (5.7–30.2) | 1.3% (0.0–3.9) | 5.9% (0.6–21.6) | 31.9% (9.1–66.3) | 18,377 (1,388–132,086) |
| Tardigrades (10) | 9.87% (1.76–18.8) | 11.8% (4.8–19.0) | 2.2% (0.52–6.4) | 15.6% (9.3–38.3) | 58.9% (36.6–78.7) | 123,043 (19,839–1,033,914) |
| Tardigrades – washed (10) | 13.1% (2.8–37.2) | 10.7% (5.4–23.7) | 1.53% (0.1–8.26) | 12.7% (4.6–27.3) | 56.9% (36.3–79.5) | 85,027 (17,658–344,379) |
| Medium (100 µl) | 1.9% (0.9–3.0) | 10.7% (3.4–13.7) | 2.8% (0.6–6.8) | 32.4% (23.9–39.0) | 52.2% (44.7–68.9) | 598,216 (217,401–2,990,684) |
| **Experiment 4** |  |  |  |  |  |  |
| Tardigrades (pool 5-50) | 7.6% (0.7–45.8) | 6.8% (2.2–18.8) | 2.8% (0.3–6.1) | 4.53% (0.0–19.3) | 71.1% (33.0–90.1) | 838,718 (36,847–15,736,667) |
| Medium (100 µl) | 3.9% (0.8–14.2) | 9.2% (3.9–28.5) | 2.8% (0.8–8.7) | 24.2% (4.9–54.5) | 58.5% (28.6–87.2) | 1,839,477 (459,048–8,347,778) |

In the two subsequent experiments, using pools of tardigrade individuals allowed us to reduce the overall contamination (Fig 2B, Table 1), and, as explained further, observe microbiota patterns specific for different species.

We have also attempted to distinguish between bacteria present in the medium, and those within tardigrades, and highlighted the latter category. However, we decided to consider both categories as tardigrade microbiota. This is because many bacteria present in tardigrade digestive tract or on the cuticle are also likely present in samples of older medium samples, and we concluded that we will not be able to distinguish reliably among the categories.

Spike-in-based template quantification, used in Experiments 3 & 4, helped us reconstruct and describe the contamination patterns. Using data on the absolute abundance of non-contaminant 16S rRNA copies (after removal of PCR and extraction contaminants), we found a strong negative correlation (Spearman’s Rho=-0.85, p<0.001) between estimated absolute abundances and the share of PCR/extraction contamination (Fig. 2C).

### Experiment 1: Broad preliminary survey

In the first Experiment, comprising individuals of 27 species and their culture media, laboratory contamination represented a large proportion of the total data (Fig. 3A), obscuring patterns among samples. After the removal of these contaminants, we obtained a long list of genotypes and OTUs among experimental samples (Table S3, Fig. 3B). Interestingly, most of these high-abundance OTUs, including OTU5 (*Solirubrobacteraceae*), OTU6 (*Undibacterium*), and OTU14 (unclassified), colonized cultures of multiple species, contradicting our expectations regarding the presence of abundant species-specific microbiota. However, there were exceptions. Specifically, the microbial community profile of individual of *Macrobiotus* cf. *recens* (PT.056) was dominated (97% of reads) by a deltaproteobacterial OTU18 classified as *Pajaroellobacter*; this bacterium, absent in all other samples in our collection, is known as a tick-vectored pathogen of cattle. Likewise, sample of *Pseudobiotus* sp. PL.318 contained 34% of OTU19 (*Alphaproteobacteria*: *Ancylobacter*, genus primarily isolated from soil), which was rare in all other samples.

**Figure 3.**
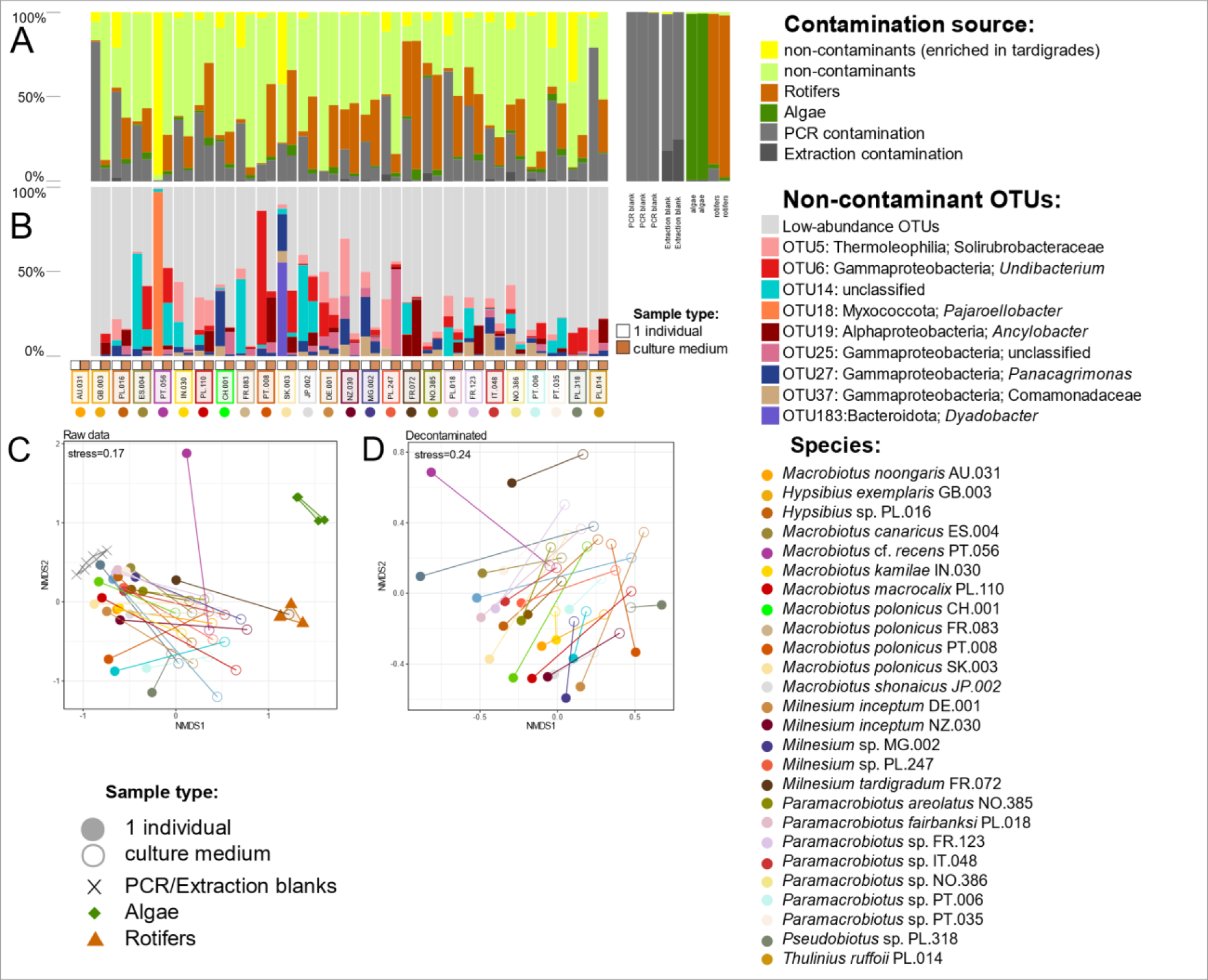
Summary of results of Experiment 1. A – classification of contamination sources; B – OTUs’ distribution in the decontaminated dataset; C – PCoA plot of raw data; D – PCoA plot of raw decontaminated data.

We also observed consistent differences among tardigrades and their culture medium samples, in the relative contribution of contaminants as well as the relative abundance of non-contaminant OTUs. The lack of within-species replicates precluded statistical comparisons, but the NMDS plot highlights substantial differences among tardigrade-medium sample pairs, especially among the NMDS2 axis (Fig. 3C). We have found significant differences between microbiota composition of tardigrade samples and the samples of culture media (PERMANOVA, *p*=0.001).

Combined, this experiment suggested that cultured tardigrades do not generally harbor abundant specialized microbiota, and to identify any such microbes it is necessary to increase signal-to-noise ratio and use biological within-species replicates.

### Experiment 2: Comparing replicate cultures

In a comparison among replicate cultures of three species, upon filtering contaminants from the microbial community profiles (Fig. 4A), the data revealed differences among the three studied species as well as among the two DNA extraction methods (Fig 4B).

**Figure 4.**
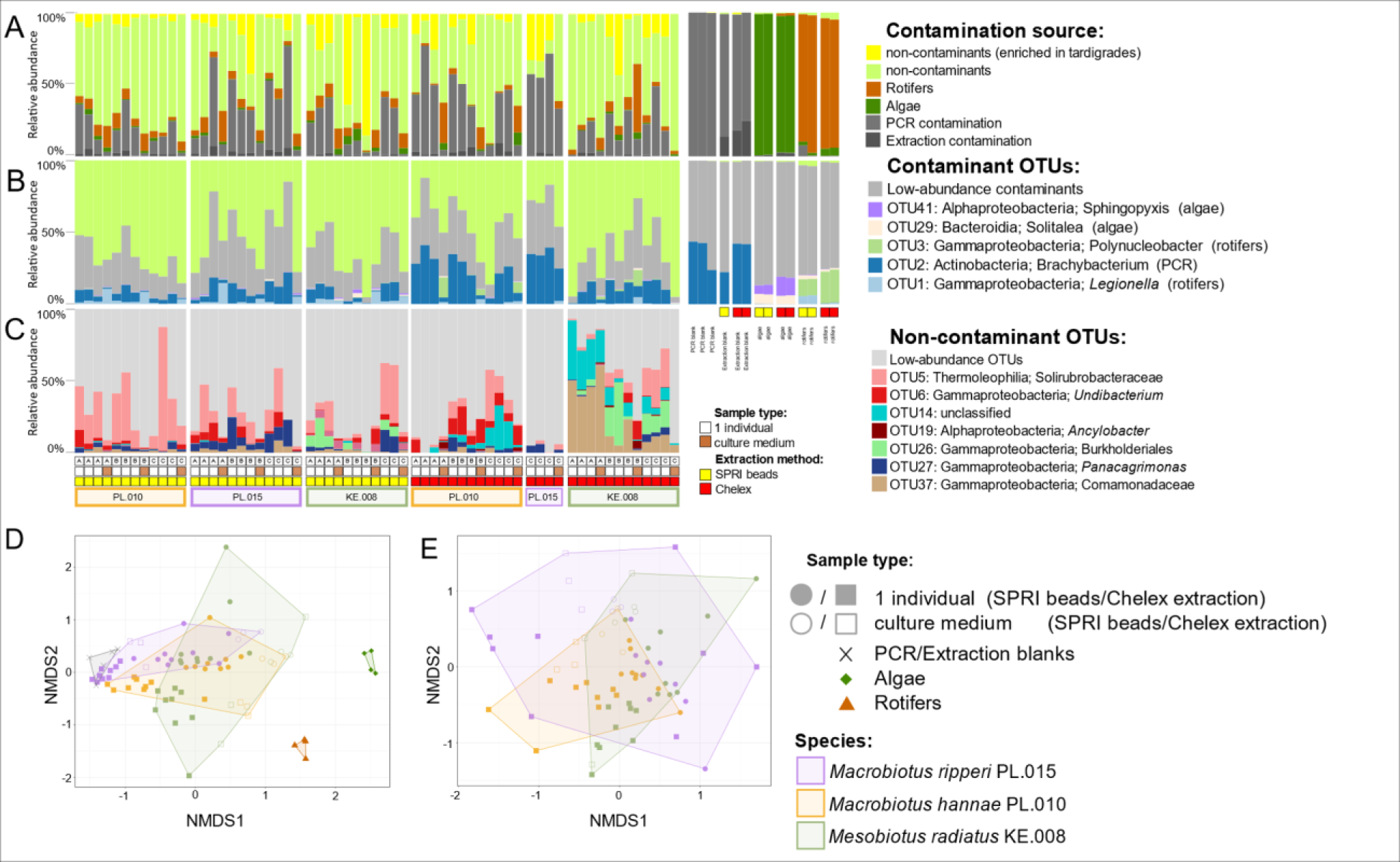
Summary of results of Experiment 2. A – classification of contamination sources; B – OTUs’ distribution in the decontaminated dataset; C – PCoA plot of raw data; D – PCoA plot of decontaminated data.

When comparing among species, we found similar OTUs across all species, but there were exceptions. The most evident exception was OTU26 (*Burkholderiaceae*) present in the individuals from all three replicated cultures of *Mesobiotus radiatus* KE.008 and, at the same time, absent in other examined cultures.

When comparing between extraction methods, we found similar taxonomic composition of dominant OTUs, but substantial differences in the relative abundances of different taxa. The most striking example of such a difference was OTU37 in *M. radiatus* KE.008 – represented by more than 25% reads in samples extracted by Chelex, and very rare (0.0%–2.17%) in samples extracted by SPRI beads. This OTU was also found in other species, with higher abundances in tardigrade samples than in culture medium.

PCoA plot of decontaminated data (Fig. 4C) shows how microbial communities vary among species and are further shaped by extraction methods. In PCoA plot, samples extracted by SPRI beads were clustered together (Figure 3 D and E), with separate groups of *M. radiatus* KE.008 distinct from *M. hannae* (PL.010) and no visible groups in *M. ripperi* (PL.015) – the species with the highest share of contaminations. The results of GLM revealed several patterns in the levels of contamination: stronger contamination in tardigrade samples compared to medium, differences between samples extracted using different methods and differences between species (Table S8).

Combined, the results of this experiment reinforced the message of the simultaneously conducted Experiment 1, showing the importance of boosting signal-to-noise ratio and using replicates, while also suggesting the superiority of the SPRI bead-based extraction method.

### Experiment 3: How to amplify the microbial signal?

In the third experiment, we tested whether pooled tardigrade individuals would provide a stronger microbial signal than single individuals, and whether we can reduce contamination by cleaning individuals with detergent. We also tested an additional extraction kit (DNeasy Blood & Tissue) as an alternative to SPRI beads, and added artificial spike-in targets as an additional control.

Similarly to the previous experiments, the contamination level was high but varied across samples and sample types (Table 1). Pooling 10 individual tardigrades per sample visibly decreased the proportions of contaminants relative to samples representing single individuals (Table 1), which was confirmed by the model (Table S9). The pooling of tardigrades also increased the share of bacteria from food sources – rotifers and algae. Overall, the contamination problem was not eliminated, as samples of culture medium had lower contaminant content than tardigrade samples. The model indicated that specimen washing did not affect the contamination content. There was a significant effect of extraction methods: samples extracted by Qiagen DNeasy Blood & Tissue kit had lower contamination levels than samples extracted by SPRI beads, but the difference was subtle – these two methods had a similar performance. The alternative extraction methods (Qiagen DNeasy Blood & Tissue kit and SPRI beads) resulted in similar microbiota profiles (PERMANOVA, p=0.08) (in contrast to Experiment 2, where Chelex resulted in a different microbial composition than SPRI beads). However, washing of individuals with detergent changed the resulting microbiota profiles (PERMANOVA, p=0.010).

Interestingly, the microbiota of pooled individuals samples was more similar to the microbiota present in culture media than the microbiota of single individuals (Figure 5E). The microbiota composition differed significantly among the three tested species (Figure 5E). Certain bacteria were more abundant in pooled samples but absent or rare in the culture medium. Such an example is OTU26 (*Burkholderiaceae*) present in *Mesobiotus radiatus* KE.008 and absent in the two other species tested in the experiment, suggesting possible species-specific association with this OTU (as indicated in Experiment 2). Apart from OTU26 in *M. radiatus* (KE.008), a visible difference between species was in the abundance of OTU9 (unclassified) which was present in *M. ripperi* (PL.015), but rare in the remaining species. However, this OTU was also found in multiple species in Experiment 1, thus it can be a part of laboratory environment. Another example was OTU11 (*Solimonadaceae*) with low abundances in *M. ripperi* (PL.015) and frequent in *M. hannae* (PL.010) as well as in *M. radiatus* (KE.008).

**Figure 5.**
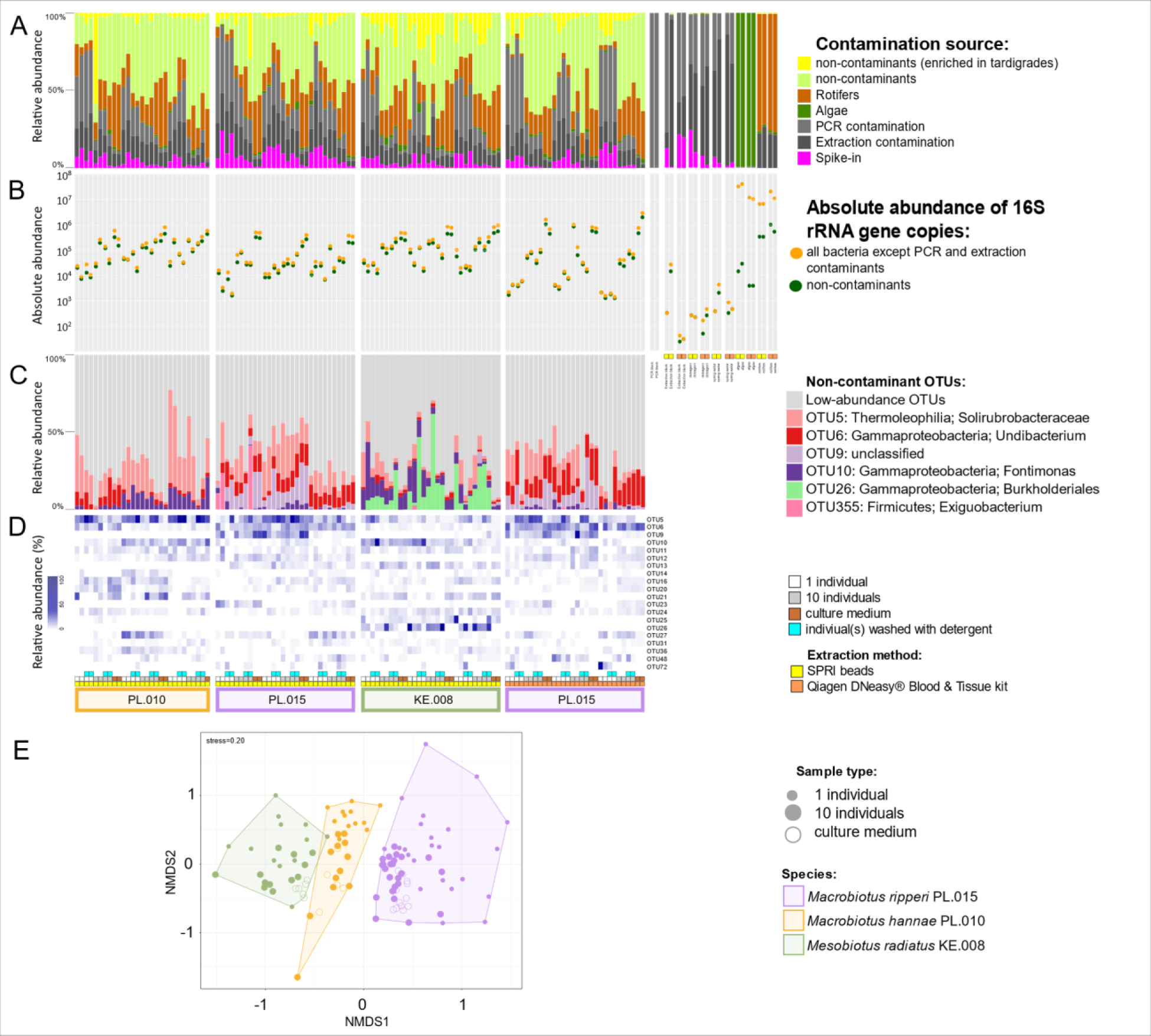
Summary of results of Experiment 3. A – relative contribution of different sources of contamination; B – estimated absolute abundance of bacterial 16S rRNA copies per sample; C – relative abundance of bacterial OTUs in the decontaminated dataset; D – the distribution and abundance of dominant bacterial OTUs across samples; E – the similarity among tardigrade and culture medium samples for different species, based on PCoA analysis of the decontaminated data.

The reads representing quantification spike-in plasmids represented a variable proportion of the libraries (range 0.03–23.95%), which varied among treatments. These values were used for the estimation of the absolute abundances of non-contaminant 16S rRNA copies across samples. Individual tardigrades contained an estimated 1,388–266,250 (median 18,377) 16S rRNA gene copies, pools of 10 tardigrades 17,658–1,033,914 (median 101,967), and samples of culture media 217,872–2,990,684 (median 589,216) copies. Tardigrade food source samples had incomparably higher abundance of bacterial 16S rRNA copies: 6,940,625–21,749,000 (median 9,216,000) for rotifers, and 10,770,000–43,015,000 (mean 23,408,000) for algae.

### Experiment 4: Final microbiota assay

We followed the strategy developed in the previous experiments to investigate the microbiota of tardigrades from 18 tardigrade cultures, in particular, attempting to maximize the number of individuals pooled per sample (Table S2, Figure 6A). This helped reduce the extent of contamination relative to the previous experiments, although it still varied among sample types and species. For most of the analyzed species it was higher in tardigrade samples compared with medium samples, with the exception of 50-individual pools of *M. ripperi* (PL.015) that were less contaminated than samples of this species culture medium.

**Figure 6.**
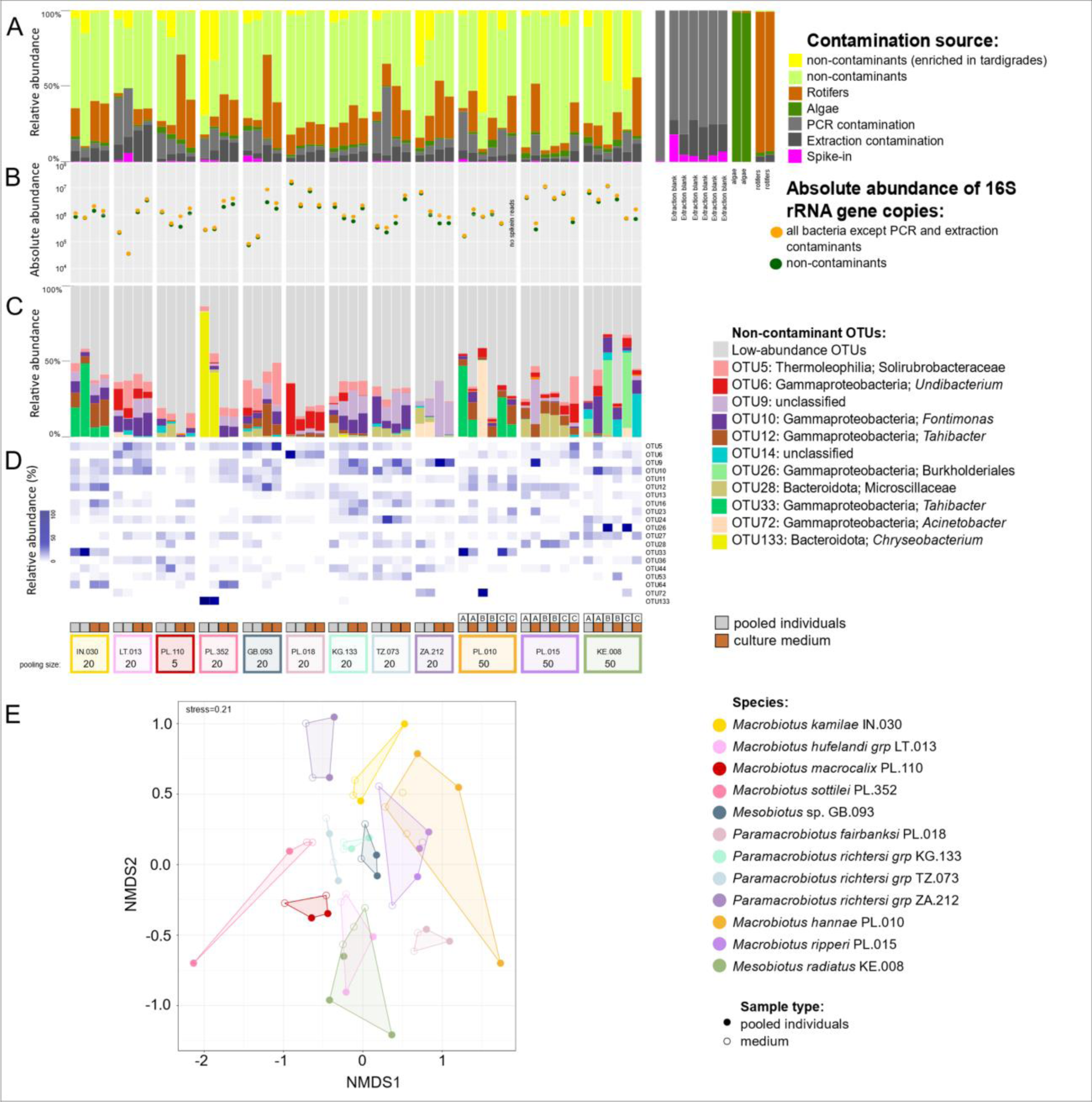
Summary of results of Experiment 4. A – classification of contamination sources; B – Absolute abundances of 16S rRNA gene copies calculated by spike-in abundance; C – OTU distribution in the decontaminated dataset; D – heatmap of most common OTUs; E – PCoA plot of decontaminated data.

The PCoA analysis revealed large differences among species in microbial community composition, as well as the similarity the microbiota of each species was similar to its culture medium (Figure 6D, 6E), particularly after decontamination. Pooling large numbers of individuals also helped identify bacterial taxa which were abundant in pooled tardigrade samples and rare in the culture medium. Examples include OTU133 (*Chryseobacterium*) present in *Macrobiotus sottilei* (PL.352), OTU26 (*Burkholderiaceae*) present in *M. radiatus* (KE.008) as well as OTU33 (*Tahibacter*) present in *M. hannae* (PL.010) and *Macrobiotus kamilae* (IN.030) (Figure 6). The estimated absolute abundances of 16S rRNA copies per sample were higher than in Experiment 3 (Figure 5B), ranging from 43,153 to 16,165,000 (mean 2,904,130) copies, and corresponding to greater numbers of individuals pooled per sample.

### Reanalysis of the data from previous studies

We reanalyzed data from the three studies on tardigrade microbiota published to date (Mioduchowska et al. 2019, Kaczmarek et al. 2020, Mioduchowska et al. 2021), together with the data from another microbiota study on freshwater mussel (*Unio crassus*) from the same lab (apparently prepared using the same protocols and reagents; Mioduchowska et al. 2020). We found that the majority of samples from across studies share multiple genotypes (zOTUs) that are among the most abundant sequences in the combined datasets (Fig. 7A). Notably, zOTUs which were found in *U. crassus* constituted, on average, 64% reads (range 3–94%) in the tardigrade libraries. As we would not generally expect such a large overlap in the genotypes among microbiota of animals as biologically dissimilar as cultured tardigrades and wild-caught mussels, we concluded that most of these zOTUs represent methodological artifacts - likely contaminants derived from laboratory reagents (noting other possibilities, including cross-contamination among samples).

**Figure 7.**
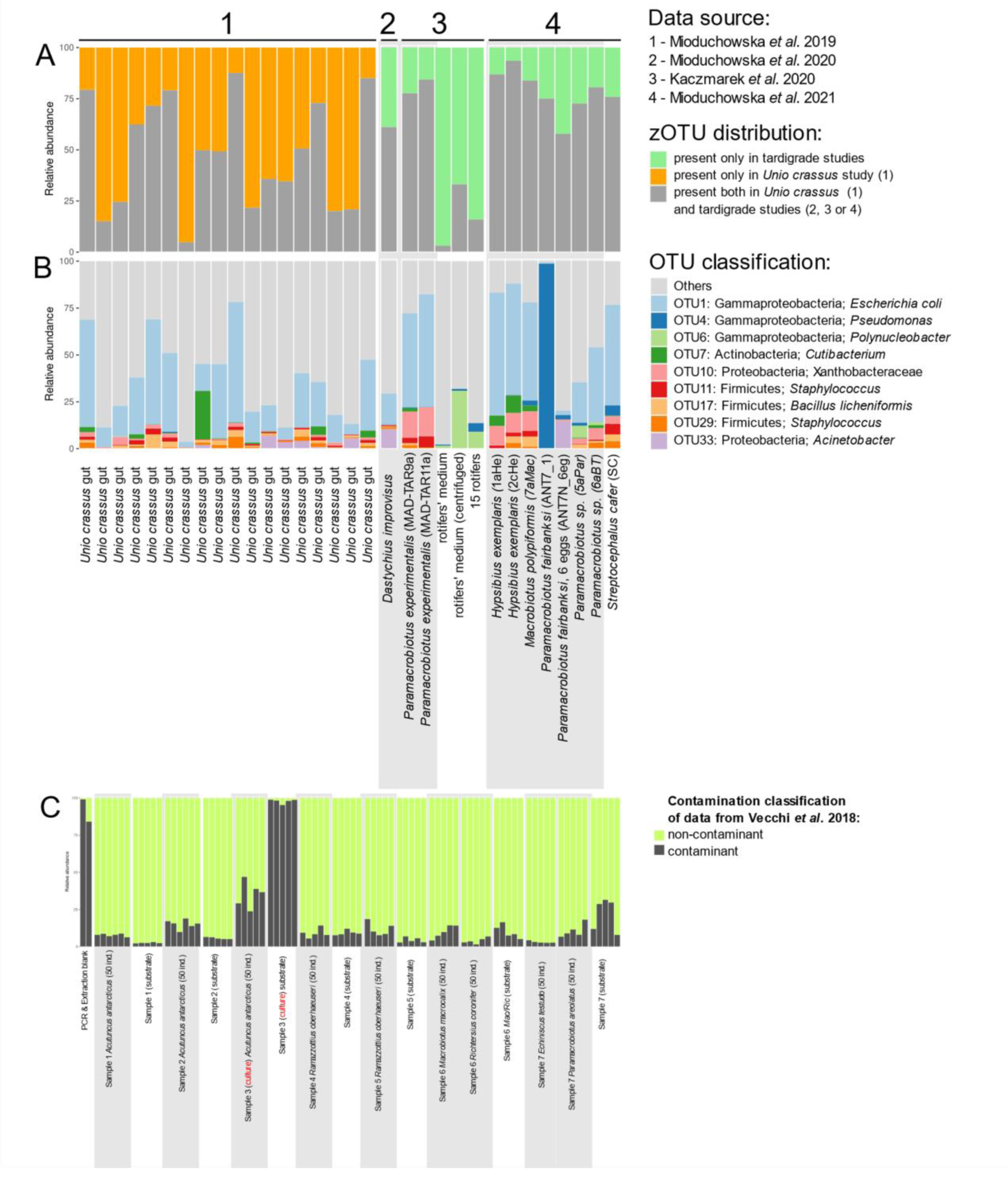
The relative contribution of putative sources of bacterial reads in the reanalysis of previously published datasets. A–B. The comparison of mussel and tardigrade microbiota data published by Mioduchowska, Kaczmarek, and colleagues. A – the summed relative abundance of zOTUs unique to mussel or tardigrade datasets, or shared among the datasets. B – the identity and relative abundance of high-abundance OTUs found in mussels as well as tardigrades (potential contaminants). C – The relative abundance of apparent reagent contaminants (zOTUs dominant in negative control samples), and other bacteria, in the study by Vecchi et al. (2018). In all panels, tardigrade samples are shaded in gray.

The inspection of taxonomic assignments of microbes identified as contaminants provided further suggestions about their nature. In particular, the most abundant contaminant was *Escherichia*, present in all samples from the four studies and constituting on average 28% (range 0–66%) of each sample (Figure 7B). The genus *Escherichia* is commonly used in the biotechnology industry for the biosynthesis of enzymes and other bioactive compounds, including molecular biology reagents (Fakruddin et al. 2013), and has been frequently reported as a contaminant in microbiota studies (Salter et al. 2014; Eisenhofer et al. 2019). Several other bacterial clades broadly distributed in this dataset (e.g., *Cutibacterium*, *Pseudomonas*) had been also listed among common contaminants. The authors of these studies did not attempt to identify reagent contaminants in their studies.

A different issue in that dataset was the presence of *Polynucleobacter* in data from Kaczmarek *et al*. 2020 (rotifer and tardigrade samples) and Mioduchowska et al. 2021 (tardigrades). The authors speculated that that bacteria could be a putative tardigrade endosymbiont. However, the high abundance of *Polynucleobacter* in rotifer samples in the present study (we used the same rotifer species *Lecane inermis* from the same source culture) suggest that it is likely a rotifer-associated microbe rather than a tardigrade symbiont.

The reanalysis of data from Vecchi et al. (2018) brought ambiguous results. The authors produced two negative control samples; however, no systematic decontamination was performed and only a few OTUs were subjectively classified as contaminants. The application of the same decontamination method as in our experimental data indicated that the overall level of contamination present in the samples was low (in most samples less than 10% of reads were classified as contaminants), with the exception of the culture medium of *Acutuncus antarcticus* (the only cultured species used in the study). Tardigrade samples from this culture had more than 25% contaminants, whereas over 95% of reads were classified as contamination in medium samples (Figure 7C). While some of the contaminants had been detected and removed by the authors (e.g., *Acidobacteria*, *Gemmata*, *Zavarzinella*), other microbial OTUs that are enriched in negative controls avoided detection as contaminants in the original study (e.g., *Pseudomonas* and *Alphaproteobacteria* OTUs).

Notably, despite the contamination issues, some of the same bacteria were present at moderately high abundance in samples from across datasets. Namely, a sequence assigned to OTU37 (*Burkholderiales*) from the present study (with 100% identity in the V4 region) was found in *Paramacrobiotus experimentalis* (Kaczmarek *et al*. 2020) (1% relative abundance) and in *Paramacrobiotus* sp. (Mioduchowska *et al*. 2021) (19–31% relative abundance). In our study, OTU including this sequence was present in 36 out of 44 analyzed tardigrade species (82%) and in 14 of them, it represented >5% of all reads after decontamination. Moreover, an identical sequence was also found in all samples in the Vecchi et al. (2018) dataset, with higher abundance in substrate samples (the highest in substrate of Sample 1: 0–7%). However, there is a possibility that this might be in fact a contaminant, as *Burkholderiales* DNA was also found at low abundance (up to 6 reads per sample) in mussels (Mioduchowska et al. 2020). Also, 8 of 18 other zOTUs assigned to this same OTU were classified as extraction or PCR contamination in the present study.

## Discussion

### Not all animals need abundant microbiota

Probably the most often asked question concerning microbiota in an unknown animal system is about the identity of the associated microbes. But the question whether there actually are any microbes to speak of is rarely considered: researchers often take the existence of abundant microbiota for granted. However, multiple studies demonstrated differences among species in the amounts of specialized, beneficial microbiota (Hammer et al. 2019). For example, while several clades of ants host abundant bacteria within their guts or cells, in about half of the surveyed species microbial amounts were at around the background contamination level (Sanders et al. 2017). Likewise, lepidopteran caterpillars were shown to harbor, on average, five orders of magnitude fewer bacteria per unit of body mass than other animals used in comparisons (Hammer et al. 2017). Our results suggest that tardigrades are also near the lower end of the spectrum of bacterial abundance. Across all experimental cultures and treatments, the contamination from reagents, food, and other sources dominated their microbial community profiles. By maximizing the number of individuals pooled into a single sample, we were able to enhance the microbial signal and observe substantial inter-species differences in microbial community composition, yet in only some cases, we observed bacterial genotypes or clades specific to a particular host species.

Tardigrades are unlikely to harbor large amounts of microbes due to their small size (generally less than 500 μm long), but we did anticipate that at least in pooled samples, we will observe a clear signal of specialized bacteria. The lack of such a signal, combined with the scarcity of genotypes enriched in tardigrades relative to culture medium, argues against the existence of specialized, abundant associations persistent in long-term laboratory cultures. The patterns contrast with our observations of abundant, specialized microbiota in some other small invertebrates. For example, in the present study, we observed abundant microbiota in rotifers, consistent across experiments. Likewise, in data that we have recently obtained for individual whiteflies (Hemiptera: Aleyrodidae, ca. 2 mm long) following the same protocols, their specialized heritable bacteria comprised ca. 99% of data on average (M. Kolasa, pers. comm.). The reasons for low bacterial amounts in tardigrades may include specific antimicrobial mechanisms, including effective immune system, production of antimicrobial compounds, behavioral adaptations, or anatomical features that limit microbial colonization. They may be combined with reduced tardigrades’ dependence on microbes for development or nutrition (Hammer et al. 2019). It is interesting to postulate, but impossible to resolve at this point, that these adaptations may correlate with tardigrades’ ability to enter cryptobiosis (Møbjerg et al. 2021). However, the lack of abundant, specialized microbiota was also observed among other microscopic animals (rotifers, cladocerans and copepods) (Eckert et al. 2021), which suggests that it may be a widespread pattern across freshwater meiofauna groups.

### Contamination as an important source of noise in microbiota studies

Microbiota studies of low biomass samples are highly prone to contamination from multiple sources, including reagents and laboratory environment. Hence, the contamination in studies of microbiota is a widespread problem, if not universally recognized (Salter et al. 2014, Davis et al. 2018). Perhaps the best-known example of contamination issues is the ongoing discussion on the human placental microbiota (Willyard 2018, Kennedy et al. 2023). Using a combination of sequencing-based approaches, Aagaard et al. (2014) detected bacteria in human placenta, earlier considered sterile, and several other authors’ works supported their claim. On the other hand, many other experiments that emphasized contamination controls failed to find evidence of microbial colonization ’*in utero*’ (eg. Lauder et al. 2016, Kuperman et al. 2020), and the discussion on whether it occurs seems to be ongoing (Kennedy et al 2023, Sharlandjieva et al. 2023).

In the field of tardigrade microbiota, an analogous groundbreaking hypothesis regarding the existence of specialized tardigrade microbiota was put forward by Vecchi et al. (2018). They detected intracellular Rickettsiales associated with samples of different tardigrade species and discussed the potential link between different reproductive modes (sexual and parthenogenetic) and intracellular bacterial infections, hypothesizing a mechanism similar to *Wolbachia*-induced reproductive manipulation in arthropods and nematodes (Kaur et al. 2021). Soon after, other authors also reported intracellular bacteria in tardigrades (Mioduchowska et al. 2019, Kaczmarek et al. 2020, Mioduchowska et al. 2021), but these conclusions were based on relatively few samples, experiments lacked replications and negative controls, and the relative abundances of putative endosymbionts were very low (much below 0.025%). Our finding that the amplicon datasets in these studies were dominated by reads almost certain to be derived from reagent contaminants, given their identity (*Escherichia*) and high prevalence in a mussell dataset processed by these authors at around the same time, showcases how the lack of stringent quality controls can lead to erroneous conclusions. Considering this, we argue that these claims of *Wolbachia* presence in tardigrades, based on few amplicon reads, need to be treated with caution until solid evidence is presented. Meanwhile, the reanalysis of the Vecchi et al. (2018) dataset, combined with diagnostic PCRs and fluorescence microscopy (Guidetti et al. 2020) failed to link reproductive modes to endosymbionts, also indicating relatively low infection rates by these microbes (9.0%–40.0%). Thus, the discussion about the diversity, specificity and roles of symbiotic bacteria in tardigrade biology remains open.

The issue of potential contamination in tardigrade microbiota studies was first mentioned by Tibbs-Cortes et al. (2022), who applied a decontamination strategy in the analysis of microbiota of tardigrade communities extracted from environmental samples. A novel component of our study is information regarding the diversity of contamination sources that may affect microbial community profiles. We identified contaminants contributed by different DNA extraction methods, originating from a PCR mastermix, as well as from different types of food. Several of the microbial genera identified as reagent-derived contaminants matched contaminants reported by previous studies (Salter et al. 2014, Eisenhofer et al. 2019). The inclusion of several types of controls enabled separation of these different contamination sources, and their effective filtering. After applying such computational filters, differences among species became much more clear (Fig. 8). Relying on relative abundance data for different microbial clades in experimental and control samples proved a powerful way of detecting and categorizing contaminants (Davis et al. 2018, Karstens et al. 2019). Still, the variation in relative abundance of genotypes across different sample types and experimental batches did complicate the conclusions about their origins. Unlike in our parallel studies of insect microbiota, where the reconstructed genotypes can often be assigned to specialized bacteria known to infect particular species and clades (e.g., Kolasa et al. 2023), here the uncertainty about the nature of different genotypes remains. We argue that even when proper controls are used, claims regarding the presence of specific bacteria in low-biomass samples need to be treated especially cautiously. Both our results and the reanalysis of the data from the previous studies argue against a major role of endosymbiotic bacteria, and highlight the lack of evidence for *Wolbachia* infection in tardigrades.

**Figure 8.**
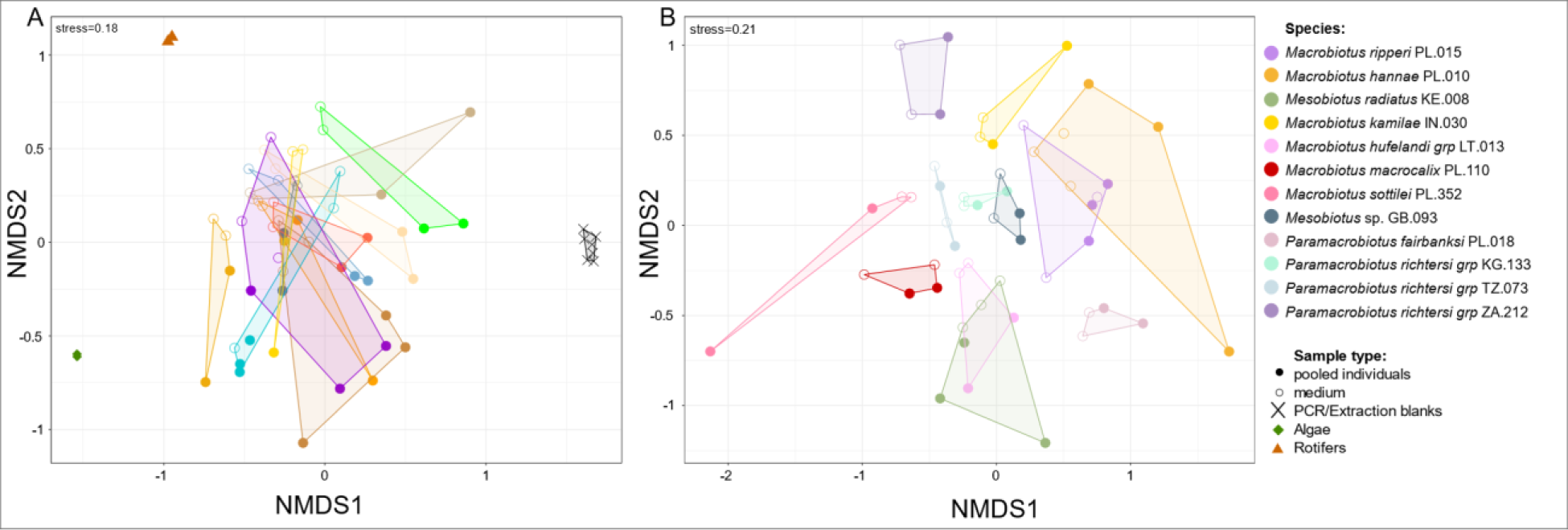
PCoA plot of samples used in Experiment 4 based on Bray-Curtis dissimilarities, calculated using A - full dataset (without decontamination), and B - decontaminated data.

### The nature, abundance, and specificity of tardigrade microbiota

Multiple invertebrate groups host abundant microbiota comprising specialized lineages reliably transmitted across generations, sometimes for tens or hundreds millions years, and playing important roles in the host biology and evolution (Moran et al. 2008). We anticipated to detect such specialized bacteria in at least some of the tardigrade cultures, inhabiting gut lumen or any internal organs, and reliably transmitted from the culture founders across generations. In our taxonomically and geographically diverse culture collection, we expected to see different specialized microbes in different cultures. We expected this category to be substantially enriched in tardigrade individuals relative to medium. On the other hand, we also anticipated that tardigrades may be colonized by less specialized bacteria, likely of environmental origin, that will thrive in lab culture environment altered by the presence of tardigrades. When thinking about the ‘core microbiota’ of tardigrades, it is important to consider these diverse origins and alternative types of associations formed by different microbes (Shade & Handelsman 2011, Risely 2020).

However, in our large collection of laboratory-cultured tardigrades, we have not found a common pattern of bacterial associations indicative of specificity. Our attempts to distinguish among bacterial associations by applying different pre-treatments – such as washing individuals in Experiment 3 – have failed to provide clear conclusions. In only a few species, we identified bacteria that are likely to be specialized tardigrade associates – including OTU26 (*Burkholderiaceae*), found in all the three experiments in samples of *Mesobiotus radiatus* KE.010, and OTU133 (*Chryseobacterium*) in *Macrobiotus sottilei* PL.352. It is plausible that some of the bacteria that are specialized but non-essential (at least in the lab environment) have been lost during years of laboratory culture. It is also likely that many species or populations other than those that we have characterized host greater amounts of specialized microbes.

Otherwise, the microbiota profiles of tardigrade specimens largely resemble environmental microbiota, with substantial overlap in microbial clades across tardigrade species and cultures. We conclude that tardigrade-associated microbiota primarily comprise bacteria of environmental origin - including those from cultures of food organisms - that are promoted in the tardigrade culture environment. At the same time, consistent differences in microbial community composition across species, observed in Experiment 4, do suggest that species alter their environment in different ways, promoting different microbes.

### Conclusions: How to study microbiota of tardigrades (and other small invertebrates from microbe-rich environments)

Studying microbiota of organisms such as tardigrades is challenging. Low biomass complicates sample handling, while low bacterial load limits signal relative to noise, making it hard to confidently separate them and accurately reconstruct host-bacterial associations (Salter et al. 2014, Knight et al. 2018). Given these challenges, you should consider if you really want to research microbiota of such systems: if you have material, resources, time, and skills to do it properly. Proving lack, low abundance, or limited specificity of microbiota is arduous and, in our experience, requires much more effort and motivation than analysis and characterization of strong and specific signal. If the authors of the present study had realized the methodological and conceptual challenges that pinpointing the tardigrade microbiota signal would require, they would have likely planned research differently.

If you decide to go ahead with the project, start from carefully defining your research questions, considering the material availability. For example, individuals extracted from natural environment or from long-term cultures will enable addressing different questions and present different challenges - such as the need to validate species identity. Ensuring that your organisms represent different species and populations, and that your samples could be considered biological replicates, would allow you to compare observed patterns in the data with biological expectations. Consider alternative means of amplifying the biological signal in your data. When working with small organisms, pooling multiple specimens into a single biological sample may be a good strategy – although that way, you lose information on variation among individuals. On the other hand, when working with wild-caught organisms, this strategy brings a risk of mixing morphologically similar species.

Consider alternative methods of generating the data. Diagnostic PCRs, marker gene amplicon sequencing, shotgun metagenomics, etc., all have their strengths, but also limitations. All these methods come with specific flavors – such as inherent differences among targeted genomic regions or sequencing platforms – and biases that you need to take into account. Regardless of the approach, it is necessary to incorporate in your experiments multiple negative controls (blanks), including DNA extraction blanks (“DNA extracted from nothing”) and PCR blanks (distilled water used for PCR reaction), and samples of any culture media and food. They will be essential for filtering and controlling microbial contamination from different sources, especially reagents. It is also worth using well-defined controls including microbial community standards, or quantification spike-in standards, as a means of quantifying and delimiting signal relative to noise. Likewise, we recommend including positive controls - reference species where you expect strong and specific microbial signal, unlikely to be obscured by contamination.

Think carefully about the bioinformatic workflows and data analysis strategy, including contaminant detection and filtering, and taxonomic levels at which sequences are classified. Standard pipelines and platforms optimized for soil or mammalian fecal microbiome data may not adequately address challenges unique to low-biomass samples; you want to verify this carefully, and consider customizing your approach. Critically interpret the data and its patterns, and compare them with biological expectations. Pay particular attention to patterns expected from reagent contamination, including the universal presence and consistent relative abundance of the same microbial genotypes across samples. On the other hand, variable genotype-level associations across individuals, populations and species suggests biological patterns.

Studying microbiota of small invertebrates from microbe-rich environments is and will remain a challenge. However, the lack, or low abundance and specificity, of microbiota may well be the biological reality in many groups of organisms, perhaps many more than commonly realized (Hammer et al 2019) - and we do want to know this when describing broad biodiversity patterns. Applying a carefully designed workflow and multiple controls should allow you to reach ground truth and uncover microbiome-related patterns in taxa such as tardigrades. Such data will provide valuable, if sometimes unexpected insights into essential aspects of biology of your study organism.

## Data availability

All data underlying this study have been deposited in NCBI, under the BioProject accession number PRJNA1066720. Details of bioinformatic workflows and scripts are available from a Github repository: https://github.com/bsurmacz/Tardigrade_microbiome.

## Acknowledgements

This project was supported by the Polish National Agency for Academic Exchange grant PPN/PPO/2018/1/00015, and Polish National Science Centre grants 2018/30/E/NZ8/00880 and 2016/22/E/NZ8/00417.

