## Supporting Information 2 for "Pinpointing the microbiota of tardigrades: what is really there?"

**Experiment 1: Broad preliminary survey**

**1. Difference in percentage of contamination between tardigrade samples and medium**

We have found significant differences between contamination levels between tardigrade and medium samples (Wilcoxon test, p<0.001).

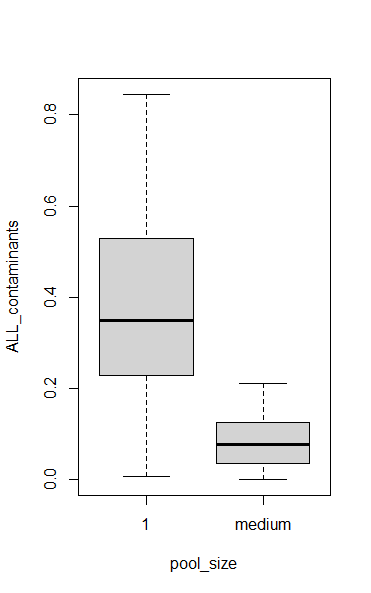

**Figure S1.** Percentage of contaminants (PCR & Extraction contamination) in libraries of tardigrades vs medium samples.

**Experiment 2: Comparing replicate cultures**

**2.1. Difference in percentage of contamination between samples, accounting for multiple factors**

We tested the effects of sample preparation methods (DNA extraction method, sample type: tardigrade/medium, species) using Generalized Linear Model using beta distribution (logit link). Culture replicate was used as a random factor.

Model 1 formula:

*Contaminants* ~ *ExtractionMethod* + *PoolSize* + *Species* + (1|*CultureReplicate*)

**Table S8. Model 1 parameters.**

|  | Estimate | Std. Error | z-value | Pr(>\|z\|) |
| --- | --- | --- | --- | --- |
| (Intercept) | -1.0274 | 0.1922 | -5.345 | 9.07e-08 *** |
| ExtractionMethod_Chelex | 0.7343 | 0.1868 | 3.930 | 8.50e-05 *** |
| SampleType_Medium | -1.6285 | 0.2379 | -6.845 | 7.62e-12 *** |
| Species_Mac.PL.015 | 1.0388 | 0.2150 | 4.831 | 1.36e-06 *** |
| Species_Meb.KE.008 | -0.5317 | 0.2346 | -2.266 | 0.0234 * |

Dispersion parameter for beta family: 7.09

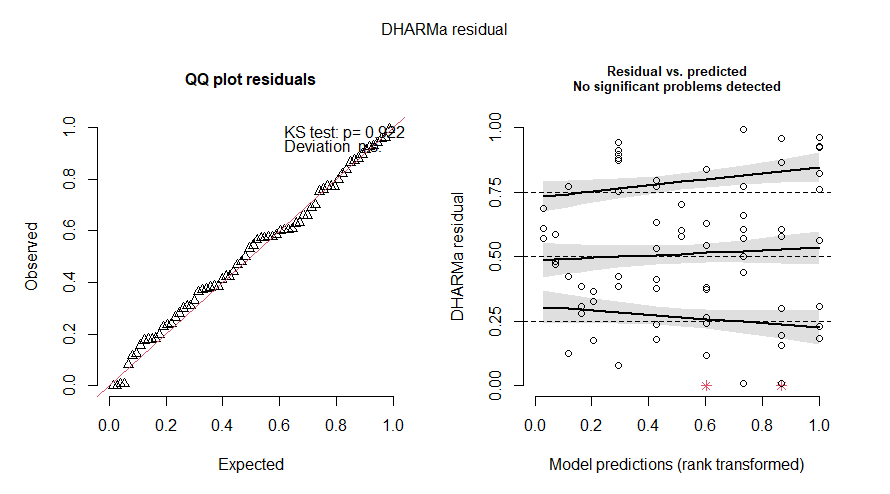

**Figure S2.** Residual plots for Model 1.

**2.2. Difference in community structure between samples, accounting for multiple factors (after decontamination)**

**Experiment 3: How to amplify the microbial signal?**

**3.1 Difference in percentage of contamination between samples, accounting for multiple factors**

We tested the effects of sample preparation methods (treatment: washed/not; sample type: tardigrade/ 10 tardigrades/medium; species) using Generalized Linear Model using beta distribution (logit link). Culture replicate was used as a random factor.

Model 2 formula:

*Contaminants* ~ *Treatment* + *SampleType* + *Species* + (1|*CultureReplicate*)

**Table S9. Model 2 parameters.**

|  | Estimate | Std. Error | z-value | Pr(>\|z\|) |
| --- | --- | --- | --- | --- |
| (Intercept) | 0.62671 | 0.15339 | 4.086 | 4.39e-05 *** |
| Species-Max.PL.015 | 0.34852 | 0.19552 | 1.783 | 0.0747 . |
| Species-Meb.KE.008 | 0.06451 | 0.19546 | 0.330 | 0.7414 |
| SampleType-10individuals | -1.02024 | 0.08901 | -11.463 | < 2e-16 *** |
| SampleType-medium | -0.93083 | 0.11864 | -7.846 | 4.31e-15 *** |
| Treatment-washed | -0.05712 | 0.08869 | -0.644 | 0.5196 |
| DNAExtraction-Qiagen | -0.26687 | 0.11123 | -2.399 | 0.0164 * |

Dispersion parameter for beta family: 20.8

Standard deviation of random effect: 0.19

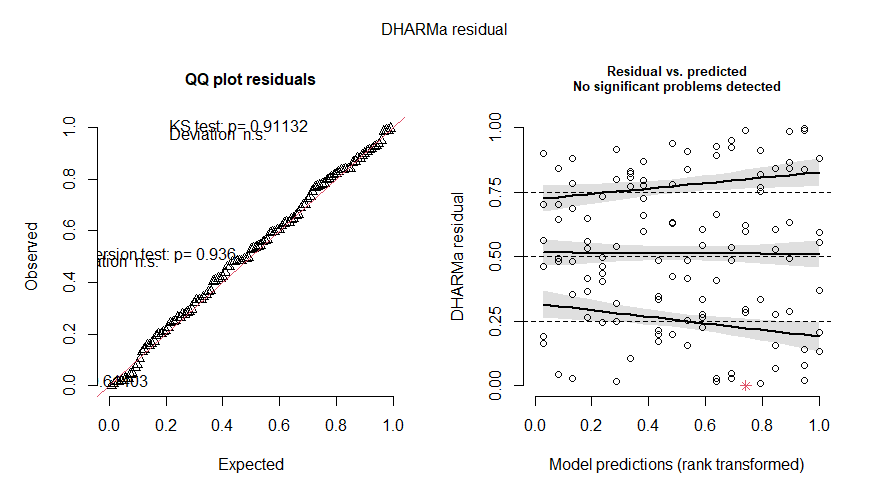

**Figure S3.** Residual plots for Model 2.
